# Cognitive performance in children and adolescents at high-risk for obsessive-compulsive disorder

**DOI:** 10.1101/2020.02.13.946061

**Authors:** Elisa Teixeira Bernardes, Carolina Cappi, Marina de Marco e Souza, Marcelo Queiroz Hoexter, Priscila Chacon, Guaraci Requena, Euripedes Constantino Miguel, Roseli Gedanke Shavitt, Guilherme Vanoni Polanczyk, Marcelo Camargo Batistuzzo

**Affiliations:** Departamento de Psiquiatria, Hospital das Clinicas HCFMUSP, Faculdade de Medicina, Universidade de Sao Paulo. R. Dr Ovidio Pires de Campos, 875, Sao Paulo – SP, Brasil; Departamento de Estatística, Instituto de Matemática e Estatística, Universidade de São Paulo. R. do Matão, 1010, São Paulo - SP, Brasil

**Keywords:** Obsessive-compulsive symptoms, high-risk, obsessive-compulsive disorder, first degree relatives, neuropsychological assessment, cognitive functions

## Abstract

**Background:** Cognitive performance has been studied in subclinical obsessive-compulsive (OC) adults and in adults relatives of OCD patients. Meanwhile, no study has been conducted with children under the same conditions. Across a sample with 49 participants, we investigated whether children and adolescents at high-risk (HR) for OCD (n=18) would present an impaired performance on neurocognitive domains compared to healthy controls (HC, n=31), especially in visuoconstructive ability, executive functions, and intellectual efficiency, functions previously associated with OCD.

**Methods:** For the HR group, we considered the first-degree children relatives of patients with OCD that present obsessive-compulsive symptoms (OCS), but do not meet diagnostic criteria for OCD. Child psychiatric diagnoses were assessed by the Kiddie Schedule for Affective Disorders and Schizophrenia (K-SADS-PL) by experienced and trained psychologists and OCS were measured by Yale-Brown Obsessive-Compulsive Scale (Y-BOCS).

**Results:** Although we did not find differences in the domains previously hypothesized, a Multivariate Analysis of Variance (MANOVA) revealed impairments in processing speed in HR group (*p* =0.019; F=3.115), and a *t-*test identified a higher IQ discrepancy in HR group when compared to HC (*p* =0.030; *t* =-2.239), and a discrepancy in verbal and non-verbal abilities as in the memory and working memory tasks.

**Conclusions:** Our results suggest that differences in motor and processing speed and in IQ discrepancy are already present and were identified in a non-clinical sample of HR subjects. Further studies should investigate neurocognitive domains as possible predictors of pediatric OCD.

## Background

Obsessive-compulsive disorder (OCD) is a neuropsychiatric disorder with a lifetime prevalence of 1.5-2.5% [1, 2], which presents its onset typically during adolescence or young adulthood [2]. It is 3 to 12 times more frequent in first-degree relatives during childhood and adolescence, and five times more frequent in adulthood [3, 4]. A Brazilian study identified that subclinical obsessive-compulsive (OC) subjects are common in first-degree relatives of individuals with OCD and are associated with lower socioeconomic status, coercive behaviors, and proband contamination/cleaning symptoms [5]. As well as the higher prevalence rate of OCD in first-degree relatives [6, 7], these family members may also exhibit a cognitive profile similar to that presented by OCD patients [8–11].

Neuropsychological studies in adults with OCD have reported impairments in visuospatial abilities, executive functions, verbal memory, verbal fluency and attention [12, 13] while in children the main impairments related are visual memory, visual organization, processing speed, cognitive flexibility and planning[14–16], although the unique meta-analysis in pediatric populations could not find any positive result [17]. In addition, adult studies with non-symptomatic first-degree relatives of OCD patients have identified cognitive deficits, particularly in inhibitory control [8, 18], decision making [9, 10], long-term verbal and visual memory [19], planning [20], working memory, verbal fluency and motor speed [11]. Regarding Intelligence Quotient (IQ), performance differences were identified in adults, in a recent meta-analysis that pointed not only an IQ difference between patients with OCD and healthy controls but also a larger IQ discrepancy between verbal and performance IQ in the OCD group [21]. Finally, Zhang et al. (2015) evaluated the cognitive performance of adults with early-onset OCD and identified impaired inhibitory control and cognitive flexibility in both patients with OCD and their siblings when compared to controls [22]. In view of possible cognitive differences between adult and pediatric OCD [12–16], it is important to evaluate the children’s first-degree relatives, which could offer a new contribution to the OCD literature.

Furthermore, considering the notion of OCD as an ongoing condition, more than a present-or-absent categorical diagnosis (i.é., a spectrum), studies have investigated cognitive performance in subclinical OC sample and found common neuropsychological mechanisms between these subjects and patients with OCD [23, 24]. When comparing cognitive performance between adults with OCD and a healthy control group, Bédard et al. (2009) identified that cognitive impairments occurred regardless of the severity of OCD [25]. Also in an adult sample, Abramovitch et al. (2015) investigated performance differences in a Go-NoGo task between a sample with subclinical OC and a control group, finding a worse performance in response inhibition and sustained attention, and concluding that subclinical OC may be associated with a moderate degree of deficient performance in these abilities [24]. Although the literature exhibits studies about the clinical and cognitive performance of adults with subclinical OC [23–25], no study has investigated the cognitive profiles during childhood with subclinical OC or in children first-degree relatives of OCD patients.

Our goal was to investigate the cognitive profile in children at high-risk (HR) for OCD and to compare it with healthy controls (HC). Here, we defined HR as first-degree relatives of patients with OCD that present obsessive-compulsive symptoms (OCS), but do not meet diagnostic criteria for OCD. We expected that children at HR for OCD would reveal some degree of cognitive impairment when compared to controls. Therefore, based on the literature on subclinical OC and family studies [8, 11, 23, 25], we hypothesize that children at HR would present a worse performance than controls on visuospatial skills, especially visuospatial memory, and executive functions (EF). Furthermore, given the recent results of the above-mentioned meta-analysis about IQ in patients with OCD [21], as a secondary objective of this study, we conducted a specific analysis evaluating IQ differences and examining verbal and non-verbal tasks. We also hypothesized that children at HR would present worse performance in non-verbal skills.

## Method

### Design and Subjects

This was a cross-sectional study with 18 children and adolescents at HR for OCD and 31 HC recruited from the community since 2011 to 2013 - Groups presented similar age and IQ and did not differ in terms of education level, handedness, and puberty development The group at HR for OCD required: 1) to be a first-degree relative (sibling or offspring) of a subject with OCD; 2) to present OCS; 3) do not meet criteria for OCD diagnosis, as evaluated by a clinician; 4) not having searched for treatment previously; 5) age between 7-18 years. Exclusion criteria were: 1) history of head injury; 2) history of substance abuse; 3) presence of intellectual deficiency or any other psychiatric diagnosis according to DSM-IV; 4) the presence of any neurological condition; 5) pregnancy or lactation. The HC group was recruited through media ads and active search at private and public schools. The inclusion and exclusion criteria were the same as the HR group, except for being a first-degree relative of a subject with OCD and present OCS. The OCD patients, first degree relatives of HR subjects, were diagnosed by professionals experienced in OCD treatment. For children, the instrument utilized was Kiddie Schedule for Affective Disorders and Schizophrenia (K-SADS-PL; [26]) and for adults, the interview was realized with the Structured Clinical Interview for DSM-IV [27]. A previous study, conducted by Chacon et al (2018) [5] assessed a similar HR group, and both studies were originated in the project Sequential Multiple Assessment Randomized Trial (SMART), coordinated in Institute of Psychiatry of the Clinical Hospital (IPq), University of Sao Paulo Medical School (FMUSP) [28].

From our data bank of OCD patients, a screening telephone call was made by an experienced psychologist to investigate if the probands had siblings or offspring, resulting in a sample of 278 possible candidates. From these candidates, 38 exhibited OCS and presented the age range (7-18 years) required as our inclusion criteria (see above). Twenty-three individuals accepted to collaborate with the research and were further evaluated with clinical scales (K-SADS-PL and Y-BOCS) at the IPq, FMUSP, but two were excluded due to the absence of OCS and three for fulfilling criteria for OCD. The remaining 18 subjects were assessed with neuropsychological instruments, which was conducted by trained psychologists and lasted 2 to 3 hours. This study was approved by the local Medical Ethics Committee of FMUSP and all participants and parents/legal guardians gave their written informed consent after having been enlightened the details of the procedure.

### Psychiatric and cognitive measures

All participants and their parents were interviewed with the K-SADS-PL administered by an experienced and trained psychologist and subjects’ OCS were measured by the Yale-Brown Obsessive-Compulsive Scale (Y-BOCS; [29]). The Petersen Puberty Scale [30] was administered to ascertain the pubertal status and the Edinburgh Handedness Inventory [31] was administered to assess handedness.

The neuropsychological battery consisted of tests that assessed the following cognitive domains: intellectual efficiency, attention, motor, and processing speed, visuoconstructive abilities, verbal and visuospatial memory, working memory, cognitive flexibility, self-monitoring, and inhibitory control. The tests and each variable used in the analysis are described in Table 1 and, detailed in the Online Resource (“Neuropsychological assessment”). In clinical and neuropsychological assessments, the instruments used were adapted or translated versions for the Brazilian Portuguese.

**Table 1.** Neuropsychological Domains and Test Employed in the Study

| Domain | Test |
| --- | --- |
| Definition | Subtest variables |
| **Intellectual functioning** | Wechsler abbreviated scale of intelligence (WASI; [1, 2]) |
| General intelligence (IQ) | Block Design; Matrix; Vocabulary; and Similarities. |
| **Attention** | Rey auditory verbal learning test (RAVLT; [3] ) |
| Endogenous processing of selecting relevant stimuli (concentrating) in the environment (e.g., objects)* | Span A; Span B<br>Trail making test (TMT) – Delis-Kaplan executive function scale (D-KEFS; [4])<br>1<sup>st</sup> condition omissions; and 4<sup>th</sup> condition sequence errors<br>Design fluency test (DFT) – D-KEFS [4]<br>DFT 1 e 2 % errors<br>Wisconsin card sorting test (WCST; [5])<br>WCST failures to maintain set.<br>Go/NoGo [6]<br>Go/NoGo omissions. |
| **Motor and processing speed** | Color-word interference test (CWIT) - D-KEFS [4]<br>Color naming time (CWIT 1); and word reading time (CWIT 2)<br>Grooved pegboard test [7]<br>Dominant hand time; and non-dominant hand time<br>TMT – D-KEFS [4]<br>5<sup>th</sup> condition time |
| **Visuoconstructive abilities** | Block Design test - WASI [1, 2] |
| Coordination of fine motor skills with spatial abilities | Rey-Osterrieth complex figure (ROCF; [8] )<br>Copy total score |
| **Visuospatial memory** | Corsi block-tapping test (CBTT) - Wechsler Memory Scale (WMS-R; [9]) |
| Memory for visual and spatial information | Forward hits<br>ROCF [10]<br>Immediate recall; and Delayed recall |
| **Verbal memory** | Digit span test (DST) - Wechsler Intelligence Scale for Children (WISC-III; [11] ) |
| Memory for verbal information | Forward hits<br>Rey auditory verbal learning test (RAVLT;[3]) |
|  | Immediate recall; and delayed recall |
| **Working memory** | DST – WISC-III [11] |
| Ability to retain information and perform mental operations from them | Backward hits<br>CBTT – WMS-R[9]<br>Backward hits |
| **Cognitive flexibility** | WCST [5] |
| Ability to change the perspectives, thinking of new possibilities for solving a problem | Perseverative errors; and categories<br>DFT – D-KEFS [4]<br>DFT 3% perseverative responses<br>Brixton [12]<br>Brixton hits<br>TMT – D-KEFS [4]<br>4<sup>th</sup> - 5<sup>th</sup> condition time difference |
| **Inhibitory control** | Go/NoGo – commission errors |
| Ability to resist an inclination to perform an action and, opting for a more convenient one | CWIT – D-KEFS [4]<br>CWIT 3 errors; CWIT 4 errors; and CWIT 3-1 time difference |

\* Attention is not a unitary system – it refers to several different capacities of how the organism comes receptive to stimuli and the ability to maintain concentration focused in stimuli over time (sustained attention). WASI – Wechsler abbreviated scale of intelligence; TMT – Trail making test; D-KEFS – Delis-Kaplan executive function scale, DTF – Design fluency test; WCST – Wisconsin card sorting test; CWIT – Color-word interference test; CBTT – Corsi block-tapping test; WMS – Wechsler Memory Scale; ROCF – Rey-Osterrieth complex figure; DST – Digit span test; WISC III – Wechsler Intelligence Scale for Children 3<sup>rd</sup> edition; RAVLT – Rey auditory verbal learning test.

### Data Analysis

Between-group comparisons of demographic and clinical variables were performed with independent sample *t*-test for continuous variables (e.g., IQ and age) and Chi-squared test for categorical variables (e.g., sex and handedness). Neuropsychological data were analyzed with Multivariate Analysis of Variance (MANOVA), using the group as the independent variable and the neuropsychological data as dependent variables. Variables that belonged to the same cognitive function were grouped according to the following neuropsychological domains: intellectual efficiency, attention, motor, and processing speed, visuoconstructive abilities, visuospatial memory, verbal memory, working memory, cognitive flexibility, and inhibitory control. More information about the neuropsychological domains and the variables that composed each MANOVA can be found in table 1 and detailed in the Online Resource – this approach grouped variables and intended to deal with the multiple comparisons’ problem. When a positive result was found at the MANOVA level, all its dependent variables were analyzed with independent *t*-tests to verify possible group differences – post-hoc analyses were corrected for Bonferroni multiple comparisons, and all analyses considered a two-tailed alpha level of 0.05. Statistical analyses were performed using the Predictive Analytics Software (PASW Statistics), version 18 (2009), the software R, version 3.4.1 (2017) and RStudio, version 1.0.44 (2016).

## Results

### Demographic results

Groups did not differ in terms of age, IQ, level of education, handedness, and puberty development. Data regarding age, years of education, handedness, and gender can be seen in Table 2. In terms of general intelligence, both groups presented an average IQ.

**Table 2.** Demographic and clinical characteristics of the high-risk group and healthy controls

|  | High-risk group (n=18)<br>n (%) / M (SD) | Health controls (n=31)<br>n (%) / M (SD) | p-value |
| --- | --- | --- | --- |
| Gender |  |  |  |
| Male | 12 (66%) | 18 (58%) | 0.551 <sup>a</sup> |
| Puberty Development |  |  |  |
| Age | 11.1 (2.4) | 11.7 (1.9) | 0.365 <sup>b</sup> |
| Handedness |  |  |  |
| Right | 16 (89%) | 30 (96.8%) | 0.377 <sup>a</sup> |
| Education Level |  |  |  |
| Years of Education | 6.1 (2.5) | 6.3 (2.1) | 0.743 <sup>b</sup> |
| Total IQ | 103.5 (12.2) | 105.6 (13.5) | 0.592 <sup>b</sup> |
| Y-BOCS |  |  |  |
| Total | 7.1 (5.3) | - | - |
| Obsessions | 3.6 (2.6) | - | - |
| Compulsions | 3.5 (2.9) | - | - |

<sup>1</sup> The Verbal – Performance difference is calculated by subtracting the performance IQ from the verbal IQ.
<sup>a</sup> Chi-squared Test \ <sup>b</sup> Independent t-test
Legend: M – mean; SD – standard deviation; IQ – Intelligence quotient; Y-BOCS – Yale Brown Obsessive Compulsive Scale;

### Cognitive results

Between-group analysis revealed that just the motor and processing speed MANOVA yielded a significant result (*p*=0.019; F=3.115). Results can be better visualized in Table 3 and Figure 1A. Post-hoc analyses with the independent sample *t*-test showed that no variable reached the proposed statistical significance within each MANOVA after the Bonferroni correction (Table 3).

**Table 3.** Clusters of Functions and the Respective Test Enrolled in Each MANOVA for Difference Within Performance

| Neuropsychological measure | F | Df | *p-value* -<br>MANOVA | *t*-test | *p-value*<br>independent<br>samples *t*-test |
| --- | --- | --- | --- | --- | --- |
| ESTIMATED INTELLECTUAL EFFICIENCY<sup>1</sup> | 2.085 | 4 | 0.099 |  |  |
| Vocabulary + Similarities + Block Design + Matrix Reasoning |  |  |  |  |  |
| Verbal IQ (Vocabulary + Similarities) | -- | -- | -- | -0.435 | 0.666 |
| Performance IQ (Block Design + Matrix Reasoning) | -- | -- | -- | 1.959 | 0.056 |
| IQ discrepancy (Verbal IQ - Performance IQ) | -- | -- | -- | -2.151 | 0.037* |
| ATTENTION | 0.718 | 7 | 0.657 |  |  |
| RAVLT span A +RAVLT span B + TMT 1 omissions + TMT 4<br>sequence errors + Des.Flu.1 E Des.Flu.2 - % errors + WCST failures to<br>maintain set+ Go-NoGo omissions |  |  |  |  |  |
| MOTOR AND PROCESSING SPEED | 3.115 | 5 | 0.019* |  |  |
| CWIT 1 + | -- | -- | -- | -0.832 | 0.410 |
| CWIT 2 + | -- | -- | -- | 0.674 | 0.504 |
| TMT 5 condition time + | -- | -- | -- | -2.291 | 0.030* |
| Grooved dominant hand time + | -- | -- | -- | -0.901 | 0.376 |
| Grooved non-dominant hand time | -- | -- | -- | -0.442 | 0.662 |
| VISUOCONSTRUCTIVE ABILITIES | 1.868 | 2 | 0.166 |  |  |
| ROCF total score copy + Block Design Test | -- | -- | -- |  |  |
| VISUOSPATIAL MEMORY | 2.788 | 3 | 0.052 |  |  |
| CBTT forward hits + ROCF immediate recall + ROCF delayed recall |  |  |  |  |  |
| VERBAL MEMORY | 0.679 | 3 | 0.570 |  |  |
| DST forward hits + RAVLT immediate recall +RAVLT delayed recall |  |  |  |  |  |
| WORKING MEMORY | 3.046 | 2 | 0.057 |  |  |
| CBTT backward hits + DST backward hits |  |  |  |  |  |
| COGNITIVE FLEXIBILITY | 2.362 | 5 | 0.057 |  |  |
| WCST perseverative errors + WCST categories + Des.Flu.3 %<br>perseverative errors.+ TMT 4-5 + Brixton |  |  |  |  |  |
| INHIBITORY CONTROL | 0.325 | 4 | 0.859 |  |  |
| Go-NoGo commission errors + CWIT 3 errors + CWIT 4 errors +<br>CWIT 3-1 time difference |  |  |  |  |  |

MANOVA – multivariate analysis of variance; Df – Degree of Freedom; TMT – Trail Making Test; Des.Flu. – Design Fluency Test; CWIT – Color-Word Interference Test; CWIT 1 = Color-Word Interference Test (naming condition); CWIT 2 Color-Word Interference Test (reading condition); CWIT SM errors = Color-Word Interference Test (total of self-monitored errors); CBTT – Corsi Block-Tapping Test; DST – Digit Span Test; ROCF – Rey-Osterrieth Complex figure Test; RAVLT – Rey Auditory Verbal Learning Test; WCST – Wisconsin Card Sorting Test. \* *p-value* < 0.05

**Fig 1.**
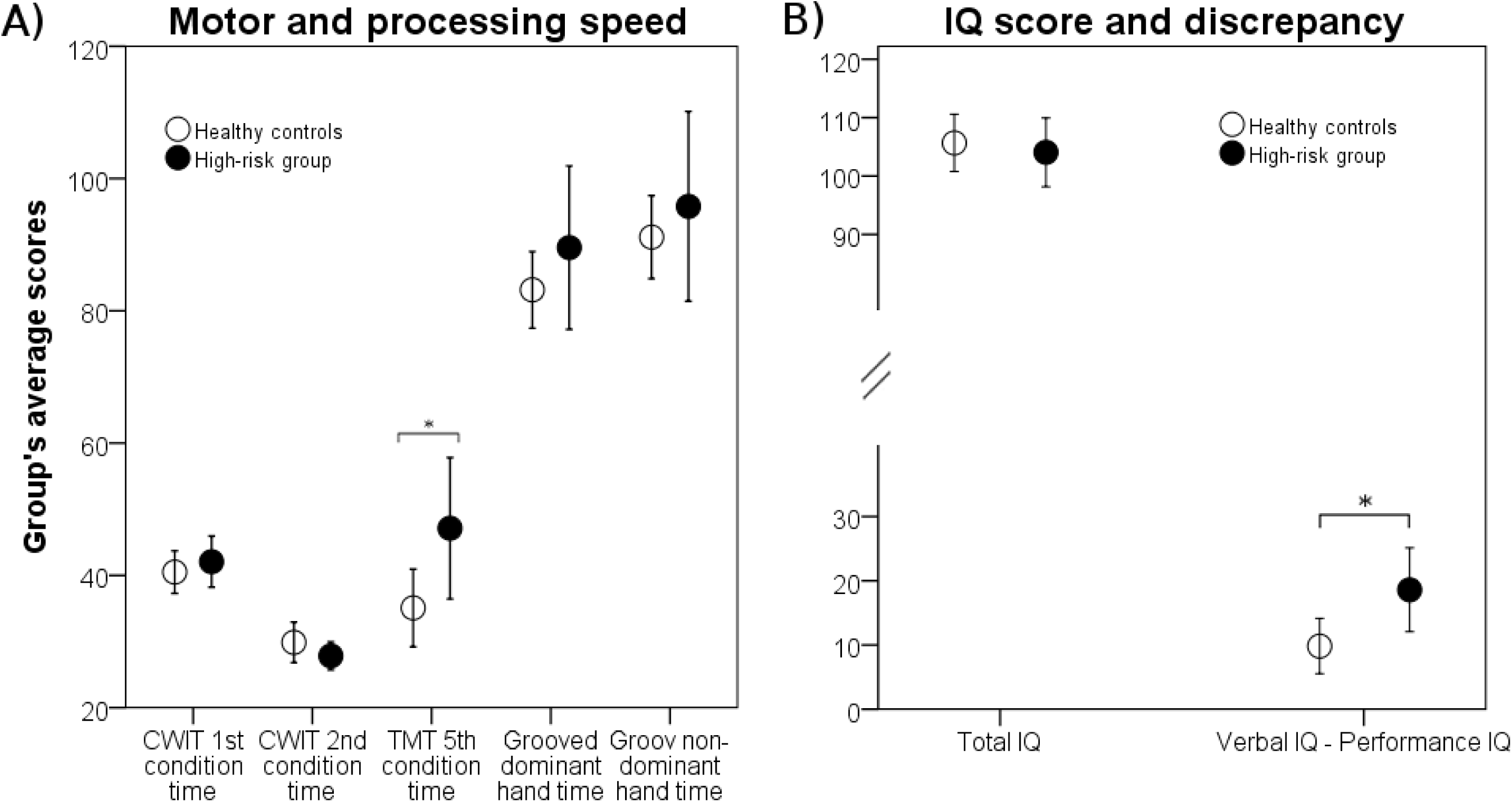
Groups’ average scores for A) the motor and processing speed MANOVA (higher punctuation means worse performance), and B) total IQ and IQ discrepancy (the difference between verbal IQ and performance IQ). Error bars means a 95% confidence interval (CI). CWIT – Color-Word Interference Test; TMT – Trail Making Test. \**p-value* < 0.05

Regarding the second objective of this study, we analyzed the IQ discrepancy (Performance Intelligence Quotient – PIQ – subtracted from the Verbal Intelligence Quotient – VIQ) between groups, and the result was significant (*p*=0.037; *t*=-2.151), indicating that the HR group exhibit a higher IQ discrepancy (Figure 1B). This IQ discrepancy was driven by a better verbal IQ and worse performance IQ in the HR group when compared to HC – separately, both IQs didn’t reach statistical significance (Online Resource, Table S1).

Finally, we wondered if this pattern observed in the IQ discrepancy would also be reflected in specific cognitive domains. In order to evaluate this hypothesis, we searched in our neuropsychological battery tests that had similar verbal and non-verbal versions, as in the case of digit span test (DST) and Corsi block-tapping task (CBTT), which offer a measure of verbal and non-verbal working memory. The HR group exhibit lower performance when compared to HC in visuospatial memory (CBTT forward, *p*=0.010, *t*=2.67) and visuospatial working memory (CBTT backward hits, *p*=0.029, *t*=2.26) (Online Resource, Table S1). On the other hand, in tests that evaluated verbal memory and verbal working memory (i.e., DST forward and DST backward), no between-group differences were identified. It is important to mention that, since we used the exploratory analysis to obtaining these results, we did not correct this analysis for multiple comparisons.

## Discussion

This study aimed to compare the cognitive performance of children at HR for OCD with HC. Our results showed differences in cognitive performance between groups in motor and processing speed: although none post-hoc analysis yielded significant results when corrected for multiple comparisons, considering a more lenient threshold (p = 0.05), we found a difference in the 5^th^ condition of the trail making test (TMT), which is a processing speed measure.

Deficits in motor speed have been identified in an adult population of non-affected OCD relatives [11] and worse performance in processing speed was also reported in the adult [21, 32] and pediatric OCD patients [14, 33]. Our study indicates that a performance difference in motor and processing speed could occur not just in pediatric OCD or adult non-affected OCD relatives, but also in non-affected children relatives with subclinical OC. Interestingly, in a review of cognitive performance in OCD, Kuelz et al. (2004) found that the information processing speed was more affected when OCD patients were medicated with selective serotonin reuptake inhibitors [34]. Our study extends these findings to a non-medicated and non-clinical sample (the HR group), highlighting the hypothesis that slower processing speed is not only related to medication usage, but also could be present in children at high-risk for the disorder without the medication usage. On this task, the subject needs to run a tracing on a dotted line, being possible that HR children present more perfectionism and obsession in performing the task, wasting more time than the HC group. Another possibility is that processing speed is a trait of the disorder [35], so that children who now have symptoms, may in future develop the disorder.

On the other hand, no significant differences were found in the other cognitive domains, including executive functions and visuospatial skills, as hypothesized. Despite the relative with the OCD diagnosis and some subclinical OC, these subjects have not searched for help or treatment because they exhibited no impairment or less of time with OCS, being considered healthy subjects. In this way, it is natural that the HR group exhibits similar performance to the control group in some cognitive functions. Especially about EF performance, studies with adults OCD subjects, OCD relatives or with subclinical OC have identified EF deficits, as in cognitive flexibility, planning and working memory [11–13, 18, 20, 24, 25]. With regard to pediatric OCD, although previous studies have identified significant results in EF performance [7, 14, 15], a recent meta-analysis could not identify any positive result [17]. These results suggest that adults and children with OCD have a different EF performance and, for a consequence, children in risk for the disorder could exhibit different results of adult relatives of OCD subjects or adults with subclinical symptoms.

Corroborating our hypothesis for the second objective from this study, results revealed that the HR group exhibited a higher IQ discrepancy than controls, with a within-group difference of 18.6 points between verbal (112.5) and performance IQ (93.9). This result is in line with a recent meta-analysis with adult OCD patients [21] that identified significant lower VIQ, PIQ, and Full-Scale Intelligence Quotient (FSIQ) scores across OCD samples when compared to controls, being the PIQ the higher difference between the groups. In consonance with the results of Abramovitch et al. (2017), we identified a VIQ-PIQ discrepancy, however with a child sample, which is a new subject-matter in the literature and possible early marker of OCD [21]. Moreover, extending these findings, the results from our last analysis point to a lower performance of the HR group in visuospatial memory and visuospatial working memory, whereas no between-group differences were identified in verbal memory or verbal working memory. Taken together, these results suggest that HR subjects may rely on verbal functions to compensate for their difficulties in visuospatial abilities. It is also possible that children with subclinical OC already exhibited the same performance established in the adult OCD literature.

As previously suggested, our two main findings – the lower performance in motor and processing speed and the IQ discrepancy – could lift another hypothesis. Processing speed is pointed as a confounding factor underlying underperformance on different domains in adults with OCD [32] and the meta-analysis with adult OCD patients conducted by Abramovitch et al (2017) suggests that PIQ can be affected by lower performance in processing speed. Therefore, it is possible that that lower processing speed plays a role in the IQ discrepancy even before the OCD establishment[21].

This study presents some limitations. First, the HR sample was relatively small and other studies with larger samples are necessary to confirm and replicate our findings. Second, it was not possible to compare our results with a group of patients with OCD. Previous adult studies compared non-affected OCD relatives or subclinical OC subjects and OCD patients and found similar results among groups. Thus, it is possible that our findings with children at HR for OCD could be replicable in pediatric OCD patients. Third, because no previous study assessed children at HR for OCD, our results were constantly compared to adult studies during the discussion – there could be differences between the adult and child populations with OCD regarding neurocognitive performance. In addition, differences in neurocognitive performance could vary in function of age, which reinforces the necessity of conducting more studies with children and adolescents at HR for OCD. Fourth, our age range was relatively large (7-18), and we know that cognitive functions, mainly EF, rely on age and the development of the prefrontal cortex [36, 37]. However, in our analysis, the subjects were paired for age, which could equally distribute this noise between both groups. Finally, our study included only first-degree relatives of subjects with OCD that present subclinical OC, not being possible to test if the results would also be applied to just one of these conditions separately (subjects with subclinical OC or first-degree relatives of OCD patients). On the other hand, this could also be seen as a study’s strength, once that no other study has included children at HR for OCD defined as being an OCD relative and presenting subclinical OC.

## Conclusion

It is possible to conclude that children at HR for OCD may exhibit a different performance in motor and processing speed and a higher IQ discrepancy when compared to HC. Once motor and processing speed was already reported in studies with pediatric OCD patients, adult first-degree relatives of OCD patients or in an adult with subclinical OC, future studies could investigate if these differences could be identified as possible cognitive marks of pediatric OCD. Also, investigating processing speed as a trait in OCD is highly relevant because impairments in this function could implicate in deficits in other cognitive functions [38]. It is also possible that the IQ discrepancy could be a cognitive mark in childhood OCD and the fact that subjects at risk for the disorder also present this characteristic early in life is of extreme relevance. Furthermore, longitudinal studies following children at HR for OCD would be of great value, allowing establishing if an impaired neurocognitive performance could present negative consequences in the future, as higher conversion rates to OCD or worse cognitive performance.

## Abbreviations

CBTT: corsi block-tapping task
DSM-IV: Diagnostic and Statistical Manual of Mental Disorder Fouth Edition
DST: digit span test
EF: executive function
FMUSP: University of Sao Paulo Medical School
FSIQ: Full-Scale Intelligence Quotient
HC: healthy control
HR: high-risk
IPq: Institute of Psychiatry of the Clinical Hospital
IQ: Intelligence Quotient
K-SADS-PL: Kiddie Schedule For Affective Disorders And Schizophrenia for School Aged-Children
MANOVA: Multivariate Analysis of Variance
OC: obsessive-compulsive
OCD: obsessive-compulsive disorder
OCS: obsessive-compulsive symptoms
PASW: Predictive Analytics Software
PIQ: Performance Intelligence Quotient
SCID-I/P: Structured Clinical Interview for DSM-IV-TR Axis I Disorders, Patient Edition
SMART: Sequential Multiple Assessment Randomized Trial
TMT: trail making test
VIQ: Verbal Intelligence Quotient
Y-BOCS: Yale-Brown Obsessive-Compulsive Scale

## Declarations

### Ethical approval and consent to participate

This study was approved by the local Medical Ethics Committee of Faculdade de Medicina, Universidade de Sao Paulo (FMUSP) and all participants and parents/legal guardians gave their written informed consent after having been enlightened the details of the procedure.

### Consent for publication

Yes

### Availability of data and material

The datasets analyzed during the current study are available from the corresponding author on reasonable request.

### Author contributions

ETB have made substantial contributions to the conception of the work; analysis, and interpretation of data; have drafted the work; have approved the submitted version (and any substantially modified version that involves the author’s contribution to the study); and have agreed both to be personally accountable for the author’s own contributions and to ensure that questions related to the accuracy or integrity of any part of the work, even ones in which the author was not personally involved, are appropriately investigated, resolved, and the resolution documented in the literature.

CC have substantively revised the work; have approved the submitted version (and any substantially modified version that involves the author’s contribution to the study); and have agreed both to be personally accountable for the author’s own contributions and to ensure that questions related to the accuracy or integrity of any part of the work, even ones in which the author was not personally involved, are appropriately investigated, resolved, and the resolution documented in the literature.

MMS have made substantial contributions to the interpretation of data; have substantively revised the work; have approved the submitted version (and any substantially modified version that involves the author’s contribution to the study); and have agreed both to be personally accountable for the author’s own contributions and to ensure that questions related to the accuracy or integrity of any part of the work, even ones in which the author was not personally involved, are appropriately investigated, resolved, and the resolution documented in the literature.

MQH have substantively revised the work; have approved the submitted version (and any substantially modified version that involves the author’s contribution to the study); and have agreed both to be personally accountable for the author’s own contributions and to ensure that questions related to the accuracy or integrity of any part of the work, even ones in which the author was not personally involved, are appropriately investigated, resolved, and the resolution documented in the literature.

PC have made substantial contributions to the conception of the work; have approved the submitted version (and any substantially modified version that involves the author’s contribution to the study); and have agreed both to be personally accountable for the author’s own contributions and to ensure that questions related to the accuracy or integrity of any part of the work, even ones in which the author was not personally involved, are appropriately investigated, resolved, and the resolution documented in the literature.

GR have made substantial contributions to the analysis of data; have approved the submitted version (and any substantially modified version that involves the author’s contribution to the study); and have agreed both to be personally accountable for the author’s own contributions and to ensure that questions related to the accuracy or integrity of any part of the work, even ones in which the author was not personally involved, are appropriately investigated, resolved, and the resolution documented in the literature.

ECM have made substantial contributions to the conception of the work; have substantively revised the work; have approved the submitted version (and any substantially modified version that involves the author’s contribution to the study); and have agreed both to be personally accountable for the author’s own contributions and to ensure that questions related to the accuracy or integrity of any part of the work, even ones in which the author was not personally involved, are appropriately investigated, resolved, and the resolution documented in the literature.

RGS have made substantial contributions to the conception of the work; have substantively revised the work; have approved the submitted version (and any substantially modified version that involves the author’s contribution to the study); and have agreed both to be personally accountable for the author’s own contributions and to ensure that questions related to the accuracy or integrity of any part of the work, even ones in which the author was not personally involved, are appropriately investigated, resolved, and the resolution documented in the literature.

GVP have made substantial contributions to the conception of the work, and interpretation of data; have substantively revised the work; have approved the submitted version (and any substantially modified version that involves the author’s contribution to the study); and have agreed both to be personally accountable for the author’s own contributions and to ensure that questions related to the accuracy or integrity of any part of the work, even ones in which the author was not personally involved, are appropriately investigated, resolved, and the resolution documented in the literature.

MCB have made substantial contributions to the conception of the work; analysis, and interpretation of data; have drafted the work; have approved the submitted version (and any substantially modified version that involves the author’s contribution to the study); and have agreed both to be personally accountable for the author’s own contributions and to ensure that questions related to the accuracy or integrity of any part of the work, even ones in which the author was not personally involved, are appropriately investigated, resolved, and the resolution documented in the literature.

## Acknowledgements

We thank Dr. Sonia Borcato and Dr. Joana Balardin for their assistance in the data collection.

## References

1. Fontenelle LF, Mendlowicz M V., Versiani M (2006) The descriptive epidemiology of obsessive-compulsive disorder. Prog. Neuro-Psychopharmacology Biol. Psychiatry 30:327–337

2. Ruscio A, Stein D, Chiu W, Kessler R (2010) The epidemiology of obsessive-compulsive disorder in the National Comorbidity Survey Replication. Mol Psychiatry 15:53–63. doi: 10.1038/mp.2008.94.

3. Nestadt G, Samuels J, Riddle M, et al (2000) A family study of obsessive-compulsive disorder. Arch Gen Psychiatry 57:358–63

4. Walitza S, Melfsen S, Jans T, et al (2011) Obsessive-compulsive disorder in children and adolescents. Dtsch Arztebl Int 108:173–9. doi: 10.3238/arztebl.2011.0173

5. Chacon P, Bernardes E, Faggian L, et al (2018) Obsessive-compulsive symptoms in children with first degree relatives diagnosed with obsessive-compulsive disorder. Rev Bras Psiquiatr. doi: 10.1590/1516-4446-2017-2321

6. Chacon P, Rosario-Campos MC, Pauls DL, et al (2007) Obsessive-compulsive symptoms in sibling pairs concordant for obsessive-compulsive disorder. Am J Med Genet Part B Neuropsychiatr Genet 144:551–555. doi: 10.1002/ajmg.b.30457

7. Pauls DL, Abramovitch A, Rauch SL, Geller DA (2014) Obsessive–compulsive disorder: an integrative genetic and neurobiological perspective. Nat Rev Neurosci 15:410–424. doi: 10.1038/nrn3746

8. Chamberlain SR, Fineberg N a, Menzies L a, et al (2007) Impaired cognitive flexibility and motor inhibition in unaffected first-degree relatives of patients with obsessive-compulsive disorder. Am J Psychiatry 164:335–8. doi: 10.1176/appi.ajp.164.2.335

9. Viswanath B, Janardhan Reddy YC, Kumar KJ, et al (2009) Cognitive endophenotypes in OCD: a study of unaffected siblings of probands with familial OCD. Prog Neuropsychopharmacol Biol Psychiatry 33:610–5. doi: 10.1016/j.pnpbp.2009.02.018

10. Lochner C, Chamberlain SR, Kidd M, et al (2016) Altered cognitive response to serotonin challenge as a candidate endophenotype for obsessive-compulsive disorder. Psychopharmacology (Berl) 233:883–91. doi: 10.1007/s00213-015-4172-y

11. Ozcan H, Ozer S, Yagcioglu S (2016) Neuropsychological, electrophysiological and neurological impairments in patients with obsessive compulsive disorder, their healthy siblings and healthy controls: Identifying potential endophenotype(s). Psychiatry Res 240:110–7. doi: 10.1016/j.psychres.2016.04.013

12. Shin NY, Lee TY, Kim E, Kwon JS (2014) Cognitive functioning in obsessive-compulsive disorder: a meta-analysis. Psychol Med 44:1121–1130. doi: 10.1017/S0033291713001803

13. Abramovitch A, Abramowitz JS, Mittelman A (2013) The neuropsychology of adult obsessive– compulsive disorder: A meta-analysis. Clin Psychol Rev 33:1163–1171. doi: 10.1016/j.cpr.2013.09.004

14. Andrés S, Boget T, Lázaro L, et al (2007) Neuropsychological Performance in Children and Adolescents with Obsessive-Compulsive Disorder and Influence of Clinical Variables. Biol Psychiatry 61:946–951. doi: 10.1016/j.biopsych.2006.07.027

15. Ornstein TJ, Arnold P, Manassis K, et al (2010) Neuropsychological performance in childhood OCD: A preliminary study. Depress Anxiety 27:372–380. doi: 10.1002/da.20638

16. Geller DA, Abramovitch A, Mittelman A, et al (2017) Neurocognitive function in paediatric obsessive-compulsive disorder. World J Biol Psychiatry 1–10. doi: 10.1080/15622975.2017.1282173

17. Abramovitch A, Abramowitz JS, Mittelman A, et al (2015) Research Review: Neuropsychological test performance in pediatric obsessive-compulsive disorder - a meta-analysis. J Child Psychol Psychiatry 56:837–847. doi: 10.1111/jcpp.12414

18. Menzies L, Achard S, Chamberlain SR, et al (2007) Neurocognitive endophenotypes of obsessive-compulsive disorder. Brain 130:3223–3236. doi: 10.1093/brain/awm205

19. Li B, Sun J-H, Li T, Yang Y-C (2012) Neuropsychological study of patients with obsessive-compulsive disorder and their parents in China: searching for potential endophenotypes. Neurosci Bull 28:475–482. doi: 10.1007/s12264-012-1262-2

20. Delorme R, Goussé V, Roy I, et al (2007) Shared executive dysfunctions in unaffected relatives of patients with autism and obsessive-compulsive disorder. Eur Psychiatry 22:32–38. doi: 10.1016/j.eurpsy.2006.05.002

21. Abramovitch A, Anholt G, Raveh-Gottfried S, et al (2017) Meta-Analysis of Intelligence Quotient (IQ) in Obsessive-Compulsive Disorder. Neuropsychol Rev. doi: 10.1007/s11065-017-9358-0

22. Zhang J, Yang X, Yang Q (2015) Neuropsychological dysfunction in adults with early-onset obsessive-compulsive disorder: the search for a cognitive endophenotype. Rev Bras Psiquiatr 37:126–132. doi: 10.1590/1516-4446-2014-1518

23. Pietrefesa AS, Evans DW (2007) Affective and neuropsychological correlates of children’s rituals and compulsive-like behaviors: continuities and discontinuities with obsessive-compulsive disorder. Brain Cogn 65:36–46. doi: 10.1016/j.bandc.2006.02.007

24. Abramovitch A, Shaham N, Levin L, et al (2015) Response inhibition in a subclinical obsessive-compulsive sample. J Behav Ther Exp Psychiatry 46:66–71. doi: 10.1016/j.jbtep.2014.09.001

25. Bédard M-J, Joyal CC, Godbout L, Chantal S (2009) Executive functions and the obsessive-compulsive disorder: on the importance of subclinical symptoms and other concomitant factors. Arch Clin Neuropsychol 24:585–98. doi: 10.1093/arclin/acp052

26. Kaufman J, Birmaher B, Brent DA, et al (2000) K-SADS-PL. J Am Acad Child Adolesc Psychiatry 39:1208. doi: 10.1097/00004583-200010000-00002

27. First MB, Spitzer RL, Miriam G, Williams JB. (2002) Structured Clinical Interview for DSM-IV-TR Axis I Disorders, Research Version, Patient Edition. (SCID-I/P). Biometrics Research, New York, USA

28. Fatori D, de Bragança Pereira CA, Asbahr FR, et al (2018) Adaptive treatment strategies for children and adolescents with Obsessive-Compulsive Disorder: A sequential multiple assignment randomized trial. J Anxiety Disord 58:42–50. doi: 10.1016/j.janxdis.2018.07.002

29. Goodman W, Rasmussen S, Price L, et al (1991) Yale-Brown Obsessive Compulsive Scale (Y-BOCS). Verhaltenstherapie 1:226–233. doi: 10.1159/000257973

30. Petersen AC, Crockett L, Richards M, Boxer A (1988) A self-report measure of pubertal status: Reliability, validity, and initial norms. J Youth Adolesc. doi: 10.1007/BF01537962

31. Oldfield RC (1971) The assessment and analysis of handedness: The Edinburgh inventory. Neuropsychologia. doi: 10.1016/0028-3932(71)90067-4

32. Burdick KE, Robinson DG, Malhotra AK, Szeszko PR (2008) Neurocognitive profile analysis in obsessive-compulsive disorder. J Int Neuropsychol Soc 14:640–645. doi: 10.1017/S1355617708080727

33. Andrés S, Lázaro L, Salamero M, et al (2008) Changes in cognitive dysfunction in children and adolescents with obsessive-compulsive disorder after treatment. J Psychiatr Res 42:507–514. doi: 10.1016/j.jpsychires.2007.04.004

34. Kuelz AK, Hohagen F, Voderholzer U (2004) Neuropsychological performance in obsessive-compulsive disorder: a critical review. Biol Psychol 65:185–236

35. Grisham JR, Anderson TM, Poulton R, et al (2009) Childhood neuropsychological deficits associated with adult obsessive-compulsive disorder. Br J Psychiatry 195:138–141. doi: 10.1192/bjp.bp.108.056812

36. Diamond A (2009) Normal Development of Prefrontal Cortex from Birth to Young Adulthood: Cognitive Functions, Anatomy, and Biochemistry. In: Principles of Frontal Lobe Function

37. Diamond A, Barnett WS, Thomas J, Munro S (2007) Preschool program improves cognitive control. Science 318:1387–8. doi: 10.1126/science.1151148

38. Sheppard LD, Vernon PA (2008) Intelligence and speed of information-processing: A review of 50 years of research. Pers. Individ. Dif.

## References

1. Wechsler D (1999) Wechsler Abbreviated Scale of Intelligence (WASI). The Psychological Corporation, San Antonio, TX

2. Trentini, C.M., Yates, D. B., & Heck VS (2014) Escala de Inteligência Wechsler Abreviada (WASI): Manual profissional. São Paulo

3. Rey A (1958) L’examen clinique en psychologie. Pressess Universitaires de France, Paris, France

4. Delis D, Kaplan E, Kramer J (2001) Delis-Kaplan executive function system (D-KEFS). Can J Sch Psychol 20:117–128. doi: 10.1177/0829573506295469

5. Heaton, R. K., Chelune, G. J., Talley, J. L., Kay, G. G., & Curtiss G (1993) Wisconsin Card Sorting Test Manual. Odessa, FL: PAR.

6. Nosek BA, Banaji MR (2001) The Go/No-Go Association Task. Soc Cogn. doi: 10.1521/soco.19.6.625.20886

7. Matthews, C. G., & Klove K (1964) Instruction manual for the Adult Neuropsychology Test Battery. Madison, Wisc.

8. Ornstein TJ, Arnold P, Manassis K, et al (2010) Neuropsychological performance in childhood OCD: A preliminary study. Depress Anxiety 27:372–380. doi: 10.1002/da.20638

9. Wechsler D. (1987) Manual for Wechsler Memory Scale - Revised. The Psychological Corporation, San Anyonio, TX

10. Osterrieth PA (1944) Le test de copie d’une figure complexe. Arch Psychol 30:206–356

11. Wechsler D (1991) Wechsler intelligence scale for children–Third edition. San Antonio, TX Psychol Corp

12. Burgess, P. W. and Shallice, T. (1997) ‘The Hayling and Brixton tests’, Bury St Edmunds, UK: Thames Valley Test Company Limited., pp. 2–4.gess PW, Shallice T (1997) The Hayling and Brixton tests. Pearson Clin 2-4

