## Supplemental material for "Cognitive performance in children and adolescents at high-risk for obsessive-compulsive disorder"

**Neuropsychological assessment**

Intellectual efficiency was assessed by the Wechsler of Abbreviated Scale of Intelligence (WASI - [1]) according to the Brazilian standardization [2], which gives the total estimated IQ and also divides scores in verbal and performance IQ. This scale is constituted by four different subtests. The two verbal subtests are 1) Vocabulary, a task in which the participant has to define the meaning of words, and 2) Similarities, an abstraction task in which the subject should establish what two different words have in common. The two non-verbal subtests are 1) Block design, a manual task that recruits visuospatial abilities, and 2) Matrix reasoning, in which the participant had to complete gaps on different designs, using logical reasoning and abstraction (but not language) by pointing the best option between five possibilities. For all subtests, a raw score was attributed to each measure that was later converted into t-scores, and, finally, to IQ.

For attention measures, the following tests were included: Rey Auditory Verbal Learning Test (RAVLT), Trail-Making Test (TMT), Design Fluency test (DFT), a computerized version of the Wisconsin Card Sorting Test (WCST; [3] and a Go/NoGo task. The RAVLT was performed with the intention of assessing verbal learning and memory. This test constitutes on a 15 word-list that are repeated 5 times – in each repetition, the subject needs to recall the maximum of words as possible. There is also an interference 15-word list and a delayed recall, after 30 minutes. The outcome variable is the sum of the words recalled after 5 repetitions and the delayed recall. Also, the span length of both word-lists was counted as an attention measure. The second test, TMT, is part of the Delis-Kaplan Executive Function Scale (D-KEFS) [4] and consists of five different conditions: a visual cancellation task and other four conditions in which the participant has to connect the dots, using numbers and/or letters. It evaluates visual attention, switched attention, motor coordination, and processing speed since the instruction is ‘to do the task as quickly as possible’. The five conditions in this test assess 1) visual scanning, 2) number sequencing, 3) letter sequencing, 4) number-letter switching and 5) motor speed. Time and errors are counted for each condition. We considered the omission errors in the 1^st^ condition and errors in following the sequence in 5^th^ condition for attention variables.

Concerning the DFT, also from the D-KEFS [4], this test assesses the ability to draw the maximum of different figures within one minute and is composed for three conditions (condition 1 – filled dots; condition 2 – empty dots only; and condition 3 switching) that assess some aspects of attention, inhibitory control (ability to ignore extraneous stimuli), cognitive flexibility and visual-perceptual speed. The variables of interest in all of this condition were the number of trials, hits, errors and repeat designs. We considered the percentage of errors in 1^st^ and 2^nd^ conditions as attention measures. Finally, two computerized tests were used to evaluate attention: the first one was a version of the Wisconsin Card Sorting Test (WCST; [3]). WCST is a classic executive functioning test in which the subject has to combine cards following a specific rule that he does not know (color, geometric form or number) and for each trial, the subject receives a feedback saying if the match is right or wrong. The WCST assesses cognitive flexibility, attention, and capability of set shifting. One measure of WCST’s is the “failures to maintain the set”, that consists in the number of times that the subject makes 5 or more consecutive correct matches but then makes an error before successfully complete the category. We considered the number of failures to maintain the set that the subject commits as an attention variable. The second test is a Go/NoGo task. On a homemade paradigm (built in E-Prime) the volunteer has to press the ‘space’ bar whenever he/she sees a letter on the computer screen, as quick as it possible (condition “Go”). The condition “NoGo” is composed of specific letters in specific colors (‘O’ in blue or ‘E’ in pink). There are a total of 96 trials: 72 “Go” and 24 “NoGo”. The task gives the quantity of error by omission (when the subject fails to press the bar), commissions (to press when should not) and hits, as well as the time to respond in the hits and commissions. The number of omissions was counted for the attention measure.

In the measure of motor and processing speed, one of the tests included was the Color and Word Interference Test (CWIT) - D-KEFS [4]. It corresponds to a Stroop task that assesses, primarily the ability to inhibit automatic responses (inhibitory control), but also evaluates cognitive flexibility, reading and processing speed. It has four conditions: color naming (condition 1), word reading (condition 2), inhibition (condition 3) and inhibition/switching (condition 4). The variables of interest in all of these conditions were time, errors and self-monitored errors (i.e., errors that the subject realizes that he has committed and corrects). For the measure of motor and processing speed, the color naming time and the wording reading time were included. Also, the Grooved Pegboard task [5] was included for motor and processing speed. In this test, the participant has to fulfill all the 25 holes with pegs in a certain order, as quick as possible. The time and the number of errors are measured for each hand separately, being the time in both conditions the variable of interest for motor and processing speed. Besides these tests, the time in the 5^th^ condition of TMT was accounted as a measure of motor and processing speed.

For visuoconstructive ability, were accounted the total score of Block design test and the copy total score in Rey-Osterrieth Complex Figure (ROCF). The test ROCF is a visuospatial task in which the subject needs to copy a complex and detailed geometrical figure, and recall it without seeing it again, after 3 and after 30 minutes. The scoring of the ROCF was based on the “36-point system” proposed by [6], and adapted by [7], that provides a raw score that ranges from 0 to 36. In the visuospatial memory, the copy total score was included in the measure of visuoconstructive abilities.

The ROCF was also included in the measure of visuospatial memory, using the total score in the immediate recall and in delayed recall. Another test accounted for this measure was the Corsi block-tapping test (CBTT), from the Wechsler Memory Scale (WMS-R; [8]), that is constituted of two conditions: a forward and a backward. In the forward condition, the test demands that the subject observes and repeats a visual sequence and in the backward condition, the subject needs to observe and respond with the backward sequence. In both conditions, the scores generated represent the raw number of hits (total score) and for visuospatial memory, the total score of the forward order was included. Concerning verbal memory, it was constituted of RAVLT and Digit Span Test (DST) from the Wechsler Intelligence Scale for Children (WISC-III; [9]). DST is a verbal test, similar to CBTT, with the difference that in both conditions (backward and forward) the subject needs to listen and repeat a numeric sequence. For composing the verbal memory, the total score of immediate and delayed recall of RAVLT and the hits in the forward condition of DST were included.

About the Executive Functions (EF), three measures were assessed: working memory, cognitive flexibility, and inhibitory control. Due to the mental manipulation of items in the backward conditions of DST and CBTT, both scores constituted the measure of working memory. Cognitive flexibility was constituted of the percentage of perseverative responses in the 3^rd^ condition of DFT, the time difference between the 4^th^ and 5^th^ in TMT, the perseverative errors and the total categories completed in WCST and, finally, for the total score in Brixton Text [10]. Brixton is a visual anticipation test, in which a blue circle goes through ten different white spaces, and the subject needs to say where the blue circle he thinks that the circle will appear in the next trial. The total score refers to the sum of hits, i.e., when a response is coherent to the logical pattern of the previous presentations. Ultimately, for inhibitory control, it was considered the commission errors in Go/NoGo test, the number of errors in condition 3 and 4 of CWIT and the time difference between the 3 and 1 conditions of CWIT.

**Table S1 – Mean, standard deviation, range and between-groups comparison of neuropsychological variables**

|  | High-risk group | | |  | Healthy controls | | |
| --- | --- | --- | --- | --- | --- | --- | --- |
|  | n = 18 | | |  | n = 31 | | |
| Neuropsychological measure | Mean | (SD) | [min-max] |  | Mean | (SD) | [min-max] |
| IQ | |  |  |  |  |  |  |
| Total (WASI) | 103.5 | (12.2) | [77-122] |  | 105.6 | (13.4) | [81-127] |
| Verbal (WASI) | 112.5 | (15.2) | [86-142] |  | 110.5 | (15.4) | [84-139] |
| Performance (WASI) | 93.9 | (9.8) | [75-110] |  | 99.7 | (10.1) | [80-124] |
| **Verbal - Performance Discrepancy¹** | **18.61** | **(13.2)** | **[-9-44]** |  | **10.8** | **(11.6)** | **[-14-33]** |
| ATTENTION | |  |  |  |  |  |  |
| RAVLT span A | 6.2 | (1.5) | [3-9] |  | 6.4 | (1.6) | [4-9] |
| RAVLT span B | 5.7 | (1.9) | [2-9] |  | 5.6 | (1.5) | [3-8] |
| TMT 1^st^ condition omissions | 0.2 | (0.7) | [0-2] |  | 0.2 | (0.4) | [0-1] |
| TMT 4^th^ condition sequence errors | 0.4 | (1.0) | [0-4] |  | 0.4 | (0.6) | [0-2] |
| DFT 1 e DFT 2 - %errors | .12 | (0.2) | [0-0.7] |  | .04 | (0.1) | [0-0.2] |
| WCST failures to maintain set | 1.1 | (1.2) | [0-4] |  | 1.2 | (1.0) | [0-3] |
| Go-NoGo omissions | 3.6 | (6.5) | [0-29] |  | 2.6 | (4.0) | [0-20] |
| MOTOR AND PROCESSING SPEED | |  |  |  |  |  |  |
| CWIT color naming time | 42.1 | (7.8) | [29-57] |  | 39.9 | (9.1) | [23-59] |
| CWIT word reading time | 27.8 | (4.3) | [21-35] |  | 28.9 | (7.4) | [18-43] |
| TMT 5^th^ condition time* | 47.1 | (21.5) | [20-89] |  | 34.0 | (14.4) | [14-74] |
| Grooved dominant hand time | 89.5 | (24.9) | [54-142] |  | 83.7 | (15.0) | [59-119] |
| Grooved non-dominant hand time | 95.8 | (28.9) | [56-166] |  | 92.4 | (19.0) | [62-148] |
| VISUOCONSTRUCTIVE ABILITIES | |  |  |  |  |  |  |
| Block Design Test | 19.0 | (9.5) | [6-34] |  | 25.6 | (12.4) | [6-48] |
| ROCF total score – copy | 28.8 | (6.5) | [13-36] |  | 30.5 | (3.7) | [22-36] |
| VISUOSPATIAL MEMORY | |  |  |  |  |  |  |
| CBTT forward hits* | 6.5 | (1.5) | [4-9] |  | 8.1 | (2.3) | [5-13] |
| ROCF immediate recall | 18.7 | (6.7) | [7.5-29] |  | 19.2 | (5.4) | [7.5-30] |
| ROCF delayed recall | 18.3 | (6.8) | [9-29] |  | 18.4 | (5.6) | [8-31] |
| VERBAL MEMORY | |  |  |  |  |  |  |
| DST forward hits | 7.5 | (1.3) | [6-10] |  | 7.0 | (2.1) | [5-10] |
| RAVLT immediate recall | 9.9 | (2.2) | [6-14] |  | 10.3 | (2.6) | [5-15] |
| RAVLT delayed recall | 9.9 | (2.3) | [7-14] |  | 10.3 | (2.9) | [4-15] |
| WORKING MEMORY | |  |  |  |  |  |  |
| CBTT backward hits | 5.9 | (1.2) | [3-8] |  | 7.0 | (1.7) | [5-12] |
| DST backward hits | 4.8 | (1.7) | [2-9] |  | 4.7 | (1.7) | [2-9] |
| COGNITIVE FLEXIBILITY | |  |  |  |  |  |  |
| WCST Perseverative error | 10.4 | (3.5) | [4-17] |  | 9.8 | (3.8) | [4-19] |
| WCST categories | 2.4 | (1.2) | [1-4] |  | 2.5 | (1.2) | [0-5] |
| DFT %Perseverative errors | .13 | (0.2) | [0-0.6) |  | .03 | (0.1) | [0-0.3] |
| TMT 4-5 | 96.2 | (86.3) | [14-292] |  | 84.3 | (50.6) | [27-279] |
| Brixton hits | 36.6 | (8.5) | [17-46] |  | 40.0 | (4.1) | [30-48] |
| INHIBITORY CONTROL | |  |  |  |  |  |  |
| Go-NoGo Commission errors | 8.8 | (3.9) | [3-18] |  | 8.5 | (3.7) | [3-16] |
| CWIT 3 errors | 2.8 | (3.5) | [0-12] |  | 2.0 | (2.8) | [0-10] |
| CWIT 4 errors | 2.0 | (2.2) | [0-8] |  | 2.5 | (3.4) | [0-14] |
| CWIT 3-1 time difference | 39.1 | (18.8) | [30-48] |  | 36.5 | (18.0) | [30-43] |

¹ The Verbal – Performance difference is calculated by subtracting the performance IQ from the verbal IQ.

WASI – Wechsler abbreviated scale of intelligence; RAVLT – Rey auditory verbal learning test; TMT – Trail making test; DFT– Design fluency test; WCST – Wisconsin card sorting test; CWIT – Color-word interference test; ROCF – Rey-Osterrieth complex figure; CBTT – Corsi block-tapping test; ; DST – Digit span test

**p-value* < 0.05

**References**

1.	Wechsler D (1999) Wechsler Abbreviated Scale of Intelligence (WASI). The Psychological Corporation, San Antonio, TX

2.	Trentini, C.M., Yates, D. B., & Heck VS (2014) Escala de Inteligência Wechsler Abreviada (WASI): Manual profissional. São Paulo

3.	Heaton, R. K., Chelune, G. J., Talley, J. L., Kay, G. G., & Curtiss G (1993) Wisconsin Card Sorting Test Manual. Odessa, FL: PAR.

4.	Delis D, Kaplan E, Kramer J (2001) Delis-Kaplan executive function system (D-KEFS). Can J Sch Psychol 20:117–128 . doi: 10.1177/0829573506295469

5.	Matthews, C. G., & Klove K (1964) Instruction manual for the Adult Neuropsychology Test Battery. Madison, Wisc.

6.	Osterrieth PA (1944) Le test de copie d’une ﬁgure complexe. Arch Psychol 30:206–356

7.	Taylor LB (1991) Scoring criteria for the ROCF. In Spreen, O., & Strauss, E. A compendium of neuropsychological tests: Administration, norms, and commentary. Oxford Univ Press

8.	Wechsler D. (1987) Manual for Wechsler Memory Scale - Revised. The Psychological Corporation, San Anyonio, TX

9.	Wechsler D (1991) Wechsler intelligence scale for children–Third edition. San Antonio, TX Psychol Corp

10.	Burgess, P. W. and Shallice, T. (1997) ‘The Hayling and Brixton tests’, Bury St Edmunds, UK: Thames Valley Test Company Limited., pp. 2–4.gess PW, Shallice T (1997) The Hayling and Brixton tests. Pearson Clin 2–4
